# MetPlast: an R package to evaluate Metabolic Plasticity using Information Theory statistical framework

**DOI:** 10.1101/2023.04.28.472208

**Authors:** Lucio D’Andrea, Leonardo Perez de Souza, Alisdair R. Fernie, Aureliano Bombarely

**Author notes:** To whom correspondence should be addressed: Prof. Bombarely, Aureliano, D’Andrea, Lucio.

## Abstract

MetPlast, is an R package that provides the functionalities necessary to analyze biological samples’ metabolic plasticity. This package takes advantage of the statistical framework provided by the Information Theory to quantify and defined metabolic plasticity parameters. Using previous implemented formula based on Shannon entropy we automatize the calculation and visualization of a set of metabolic plasticity indexes including metabolic diversity, metabolome specialization, and metabolite specialization. We use a publicly available metabolic data set on tomato domestication to demonstrate the power of the present tool to evaluate changes in metabolic plasticity parameters. Thus, MetPlast represents a new and invaluable member of the computational metabolomics toolbox, that will certainly help scientists to unveil hidden information in cell metabolomic landscape.

**Availability and implementation:** Freely available at https://github.com/danlucio86/MetPlast

## Introduction

Although arguably controversially given the true coverage afforded by current analytical techniques (Alseekh et al., 2021), the metabolome can be defined as the *complete* set of low molecular weight chemicals, named metabolites, found within a biological sample (Razzaq et al., 2019). Yet, metabolic plasticity is the ability of an organism or species to fine-tune the production of thousands of metabolites based on their proximate environment (Shaar-Moshe et al., 2019). This characteristic does not only apply at the organismal level. In fact, metabolic plasticity might also play a major role in evolutionary processes such as species adaptation, and domestication(Moghe et al., 2017).

Information theory provides a statistical framework to assess individuals (and species) metabolic plasticity(Li et al., 2020, 2016). Shannon’s entropy set the grounds to calculate different metabolic plasticity parameters such as metabolome diversity (H_j_ index), metabolome specialization – aka metabolic profile specialization -(δ_j_ index), and metabolite specialization – aka metabolic specificity of individual metabolites-(S_i_ index). Recent analytical and computational advancement has led to obtained reasonable pictures of a sample’s metabolome(Tsugawa et al., 2021). Thus, quantified MS/MS precursors and/or identified metabolites obtained in metabolomic studies can be used to estimate metabolic plasticity.

Here we present MetPlast, an R package that integrates all the calculations as well as visualization methods necessary to evaluate metabolic plasticity variations among a set of samples.

### Framework

The statistical framework to calculate the metabolic plasticity related parameters were previously described (Martinez and Reyes-Valdes, 2008; Li et al., 2016). Shannon’s entropy formula can be used to calculate a sample metabolic diversity (H_j_) considering the relative frequency P_ij_ of each metabolite (j) in each sample (i) (Fig 1):

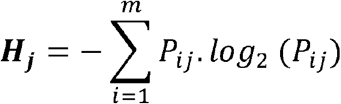

To calculate the metabolome specialization index (δ_j_ index), some intermediate calculations need to be performed. Considering the average frequency P_i_ of the ith peak among all samples, and the metabolite specialization index (S_i_), the δ_j_ index can be determine (Fig 1):

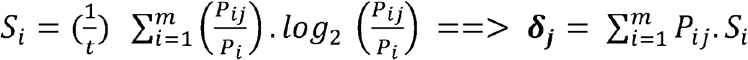

It is important to take into account that based on the fact that these functions rely on frequencies, the specialization-related parameters are not absolute quantities.

**Figure 1.**
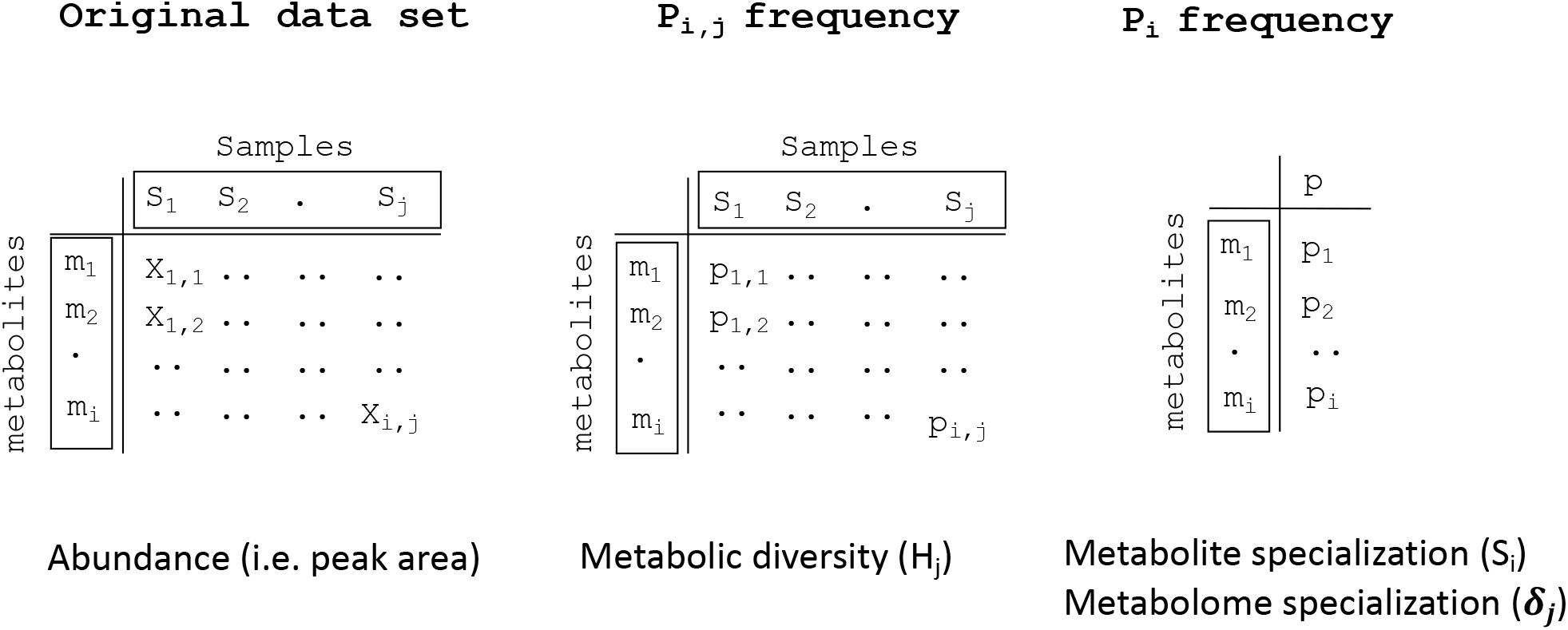
MetPlast statistical framework. MetPlast Shannon-entropy parameters (H_j_, S_i_, and δ*j*) are calculated from frequencies (p) given an data set with the abundance of metabolites 1 to i (m_1_ to m_i_) in samples from 1 to j (S_1_ to S_j_). While H_j_ is calculated from the frequency of a metabolite in a given sample (p_i,j_), S_i_ and δ*j* considers the frequency of a given metabolite (p_i_) across the whole data set.

A wide range of statistical tests can be applied to evaluate the significance difference between samples metabolic plasticity predictors. Usually, the distribution of these parameters’ values follows a normal distribution. Thus, statistical parametric comparisons tests such as ANOVA or MANOVA can be applied to evaluate statistical significance. The proper scrutinization of H_j_ differences between samples, should include the measurement of the divergence with respect to the whole average metabolome (HR_j_):

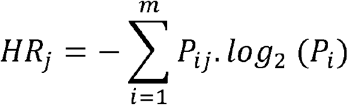

HR_j_ will be equal to or larger than the corresponding H_j_, reaching equality if and only if P_i_ = P_ij_ for all values of i. Thus, we can measure how much a sample j departs from the corresponding metabolome distribution of the whole system, by defining the Kullback– Leibler divergence (Divj):

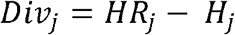

### Package structure and usage

MetPlast has 23 functions organized in 4 different categories (Table 1). The *Internal* category includes support functions. Functions included in *Individual* and *Summary* categories are intended to perform a single action towards the calculation or visualization of individual parameters. We created `Dj_index_weight()` and `Dj_index_weight_plot()` functions that evaluate the contribution of each metabolite to the δ_j_ index. Finally, *Pipeline* utilities merge two or more functions providing a quick, but complex, output that allows a complete evaluation of each parameter indicating the initial data set.

**Table 1.**
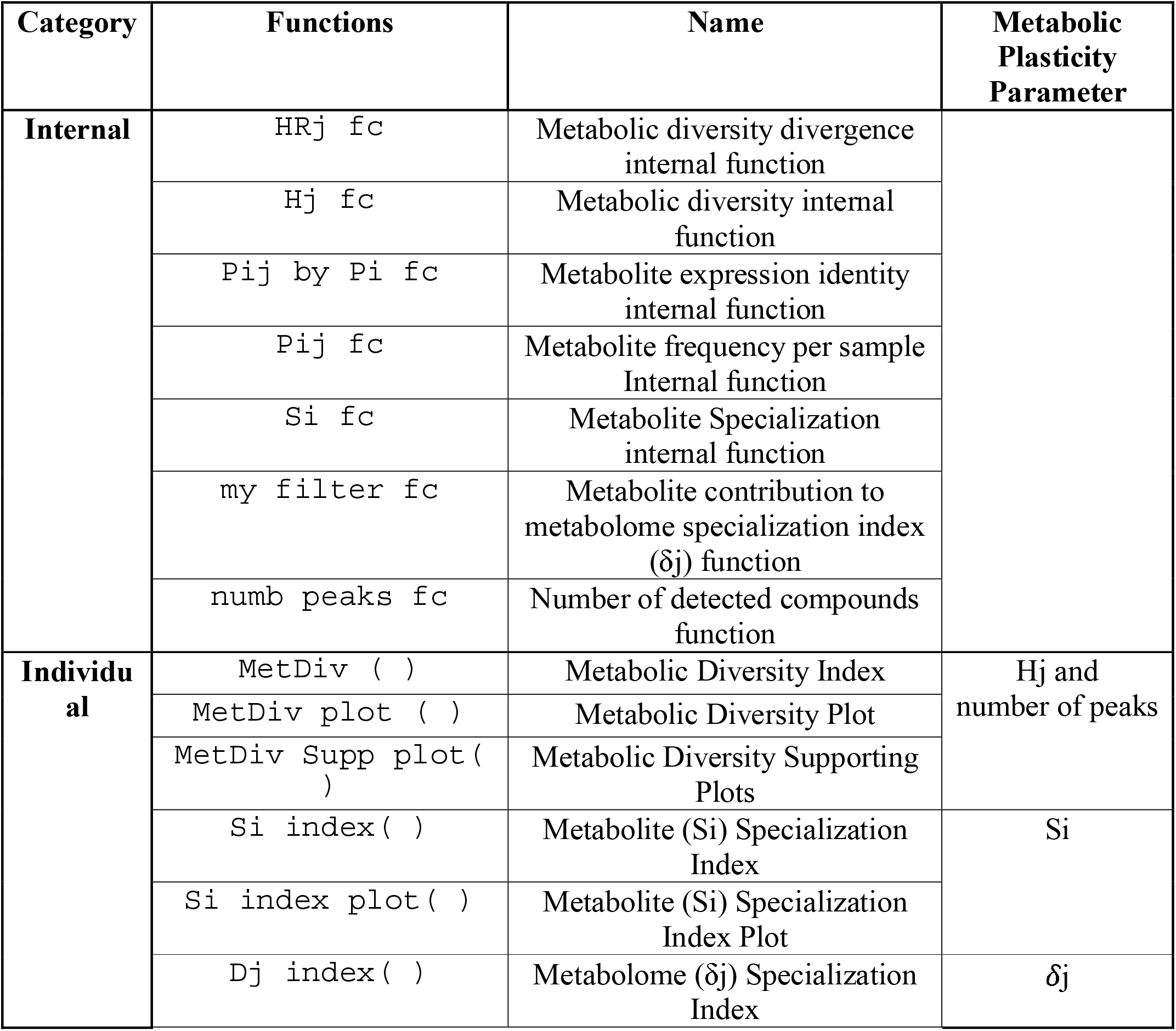
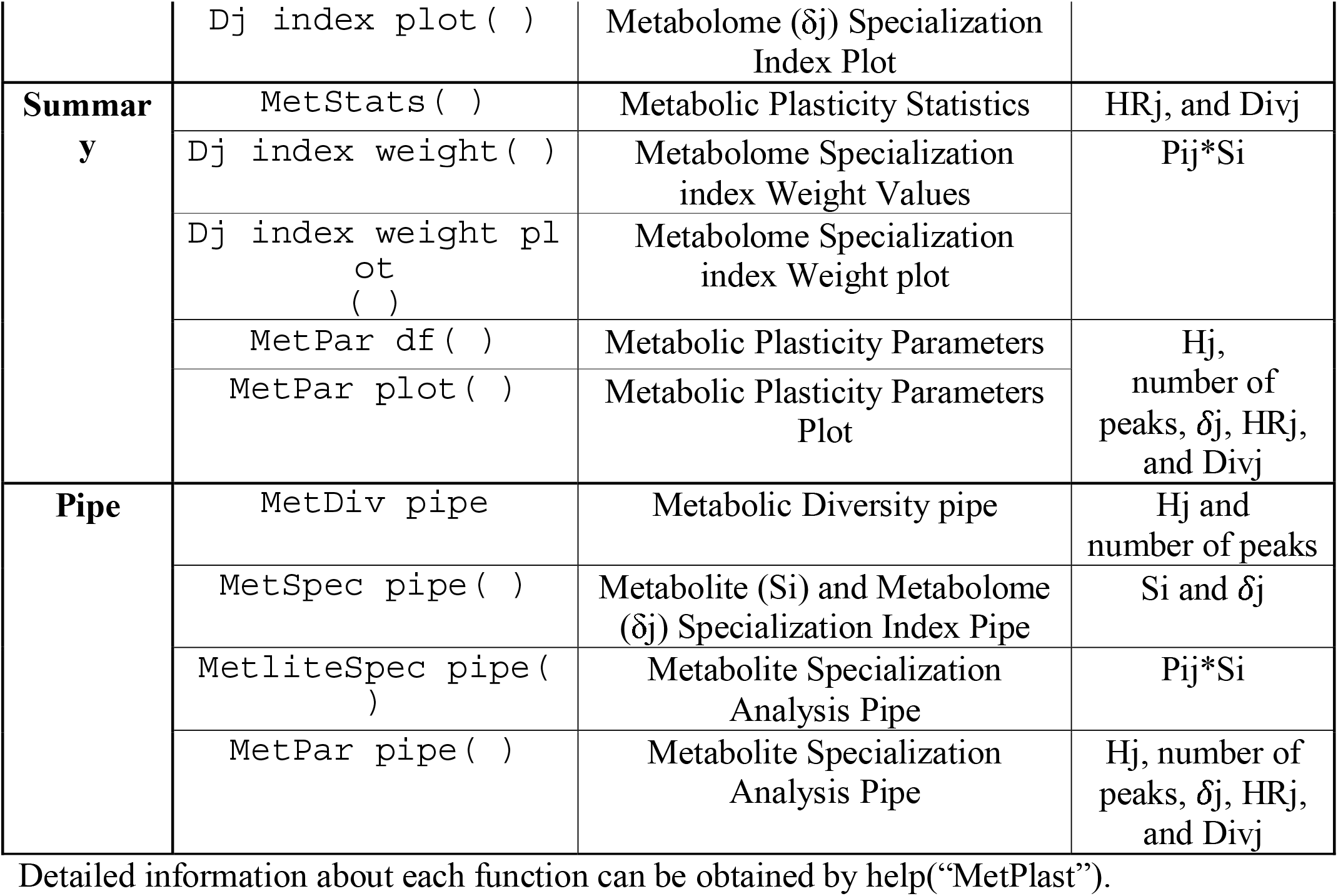
Functions available in MetPlast.R

The documentation, and usage of each function, as well as internal data sets, can be accessed using the `help (“function”).` The input data set must be a data frame with rows depicting all the metabolites/compounds detected in any of the samples ordered in columns, with the corresponding names as headers. MetPlast has been initially developed using MS data sets. Missing or negative values should be treated using either standard or more sophisticated methodologies as described in (Seekaki and Ogata, 2017).

### User case example: Tomato Domestication

MetPlast can be used to evaluate changes in the metabolic diversity during important agricultural processes such as, for example tomato domestication, using publicly available datasets (Zhu et al., 2018) (for details https://github.com/danlucio86/MetPlast). This data set includes samples from three different species, the domesticated tomato *S. lycopersicum*, and the wild relatives *S. lycopersicum var. cerasiformis*, and *S. pimpinelifolium*.

Initially, the user needs to: 1. Install the package from GitHub by the command `devtools::install_github(“danlucio86/MetPlast”)` and `library(“MetPlast”);` 2. Read the input data set – i.e. <TomDomestication.csv> - with the command `read.csv2(file=“TomDomestication.csv”,header=TRUE,row.names=“Compounds”)`; and 3. Replace missing values as described before. In general terms, we call *Data_raw* to the initial data set that might contain NA values, while we reserve the term *Data* for those data sets with no missing values. The package contains both data sets that can be easily downloaded from the Data folder.

Several analyses can be performed using MetPlast package. In order to evaluate changes in the metabolic diversity index (H_j_) during tomato domestication, we estimated the Hj using the command `MetDiv()`, and visualize using `MetDiv_plot()` (Fig 2). While Fig 2A shows the retrieved data frame with Hj values as well as the number of peaks detected in each of the samples. The evaluation of Hj per species shows that the median of the domesticated *S. lycopersicum* is lower, although not significant, than its wild relatives (Fig 2B). Possibly, the high variability observed in the Hj measurements is associated with the sampling procedure.

**Figure 2.**
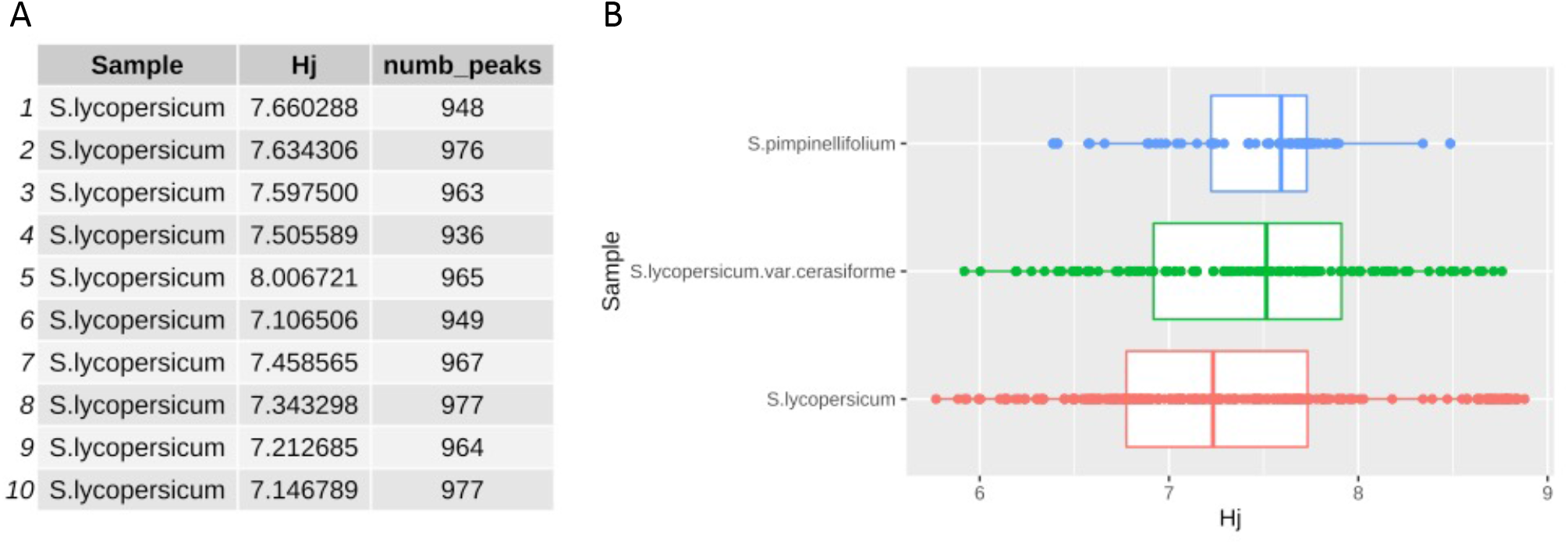
Effect of domestication in metabolic diversity (H_j_). **A**. Data frame with H_j_ values and number of peaks detected (i.e. mass features) associated with each sample under analysis. **B**. Boxplot depicting samples Hj grouped by species. Domesticated *S. lycopersicum* samples are shown in red, semi-domesticated *S. lycopersicum* var. *cerasiformis* in green and their wild relative *S. pimpinellifolium* in blue. ANOVA analysis show no statistical significance differences between the different species (p-value > 0.05).

Similarly, the specialization parameters can be estimated with the commands `Si_index()`, `Dj_index_weight()`, and `Dj_index()`, and visualize with the associated `plot()`functions (Fig 3). It is worth to mention that these parameters depend on the data set under study. The Si values associated with each metabolite is depict in Fig 3A, being the most specific, the metabolite named SlFM1980. The specialization index (Dj) is calculated by the summation of the Pij*Si. Thus, a sample Pij*Si measures the weight or impact of each metabolite in the metabolome specialization index (Dj) of a given sample (Fig 3B, C).

**Figure 3.**
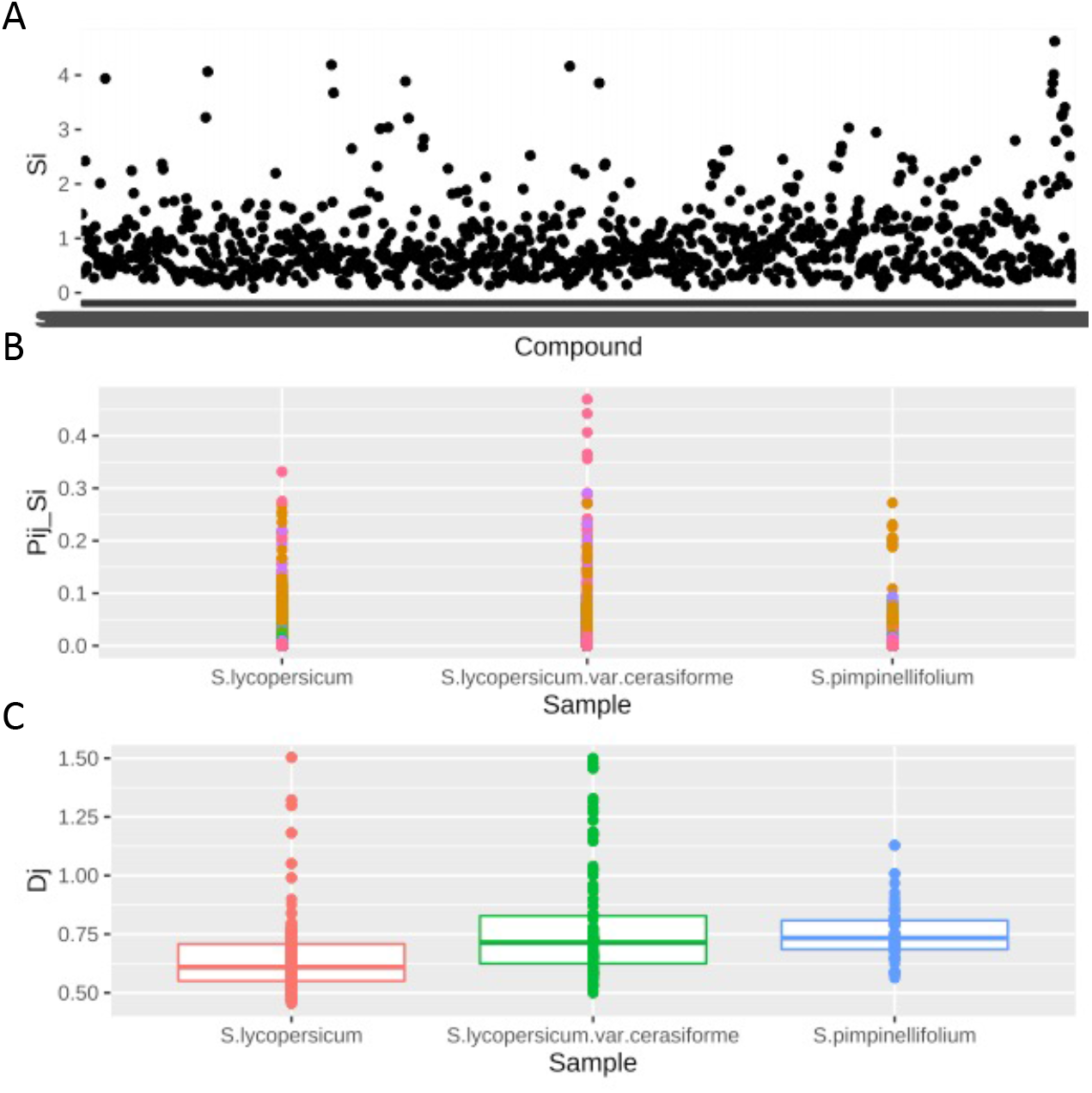
Effect of domestication in metabolite and metabolome specialization weight and index. **A**. Metabolite specilization (S_i_) has been calculated and plotted for each individual compound or metabolite. **B**. Dot plot depicting samples metabolome specialization weight, P_ij_*S*i* (Pij_Si) grouped by species. **C**. Boxplot depicting samples metabolome specialization index, δ*j* (Dj) grouped by species. Domesticated *S. lycopersicum* samples are shown in red, semi-domesticated *S. lycopersicum* var. *cerasiformis* in green and their wild relative *S. pimpinellifolium* in blue. ANOVA analysis show XXX statistical significance differences between the different species (p-value > 0.05).

Finally, specific statistics parameters, such as HR_j_ as well as Div_j_ can be calculated using the `MetStats()` command. `MetPar()` allows the user to generate an integrative data frame with all sample-specific parameters (Fig 4A). Thus, we can evaluate the specialization strategy reflected in each metabolome by plotting Div_j_ against D_j_ (Fig 4B). Thus, samples with D*j* > Div*j*, e.g. dots above the black line, are expected to have an specialization strategy that consists mainly in accumulating highly specialized metabolites, whereas samples with D*j* < Div*j* achieve their specialization by accumulating at higher or lower levels metabolites that are, on average, accumulating in the whole system.

**Figure 4.**
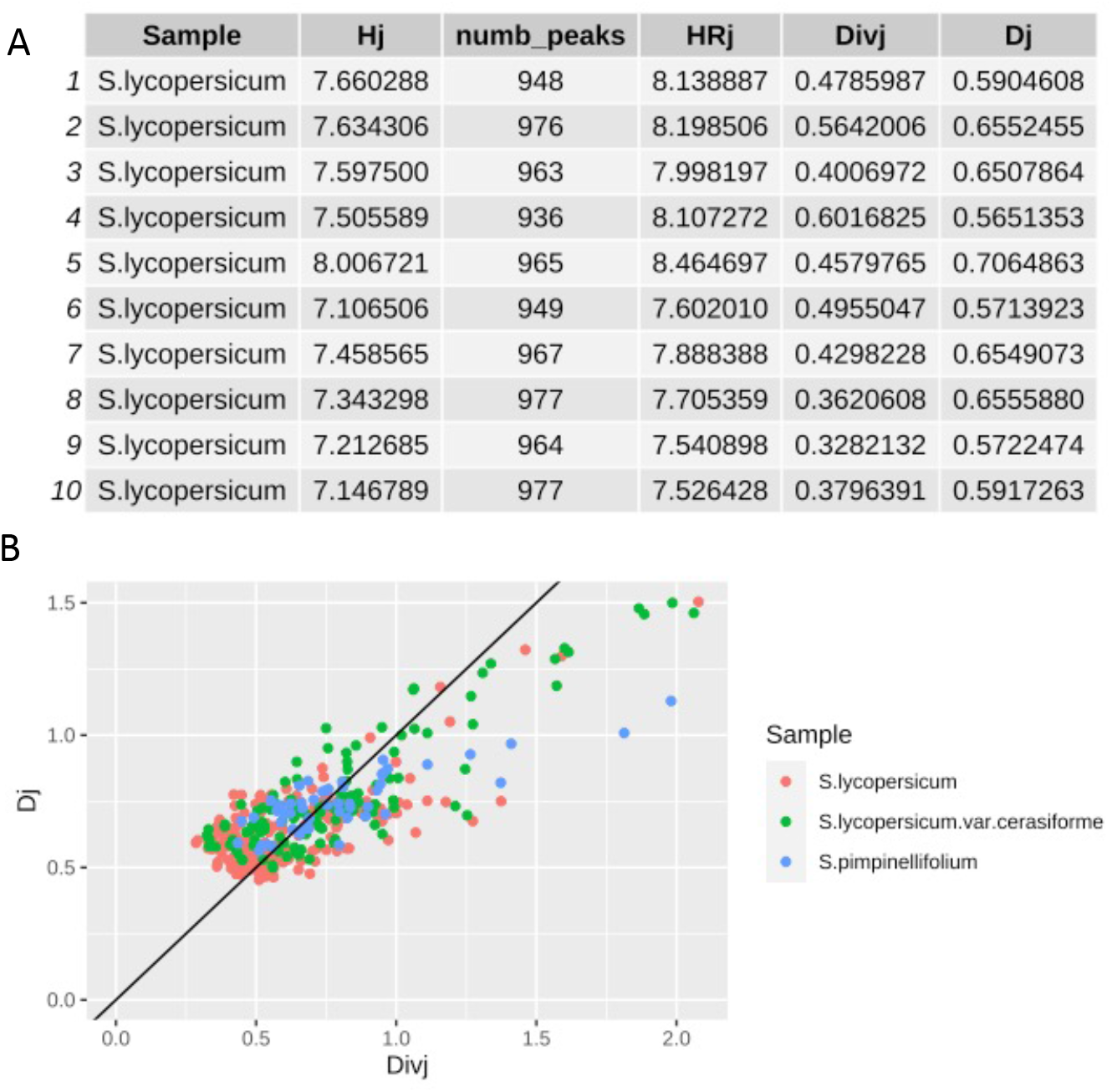
Effect of domestication in metabolic specialization strategy. **A**. Data frame integrating all the sample associated indexes and statistical indicators, e.g. metabolic diversity (Hj), number of peaks (numb_peaks), Hj divergence (HRj), and Kullback–Leibler divergence (Divj) **B**. Dot plot depicting samples Divj versus metabolome specialization index δ*j* (Dj) grouped by species. Domesticated *S. lycopersicum* samples are shown in red, semi-domesticated *S. lycopersicum* var. *cerasiformis* in green and their wild relative *S. pimpinellifolium* in blue.

Considering the different parameters that can be easily calculated using MetPlast we can extract preliminary conclusions. Our analysis on the effect of domestication on metabolome plasticity in tomato shows that there is a greater variability across samples from domesticated species. Although, this prevents a better understanding of the dynamics associated with metabolic diversity during the domestication process, it also suggests the variability is highly dependent on the sample characteristics (location, growing conditions, etc). Furthermore, on average *S. pimpinellifolium* tends to follow a metabolome specialization strategy that consists in accumulating a myriad of highly specialized metabolites. Thus, MetPlast R package helps to quickly assess a possible impact of domestication on metabolic diversity and specialization. Also allows to spot those metabolites that contributes in the process of metabolome specialization.

The simplicity of this newborn tool, which will continue to grow with the implementation of new utilities in the near future, allows users easy and rapid detailed evaluation of the dynamics associated with metabolic plasticity.

## Acknowledgement

ARF is supported by the European Union’s Horizon 2020 research and innovation programme, project PlantaSYST (SGA-CSA No. 739582 under FPA No. 664620). LD was supported by a Department of Excellence grant from MUR.

I thank to Saleh Alseekh for his expertise and assistance throughout all aspects of this study.

